# Genetic Diversity Analysis for Heat Stress Tolerance in Chili (*Capsicum annum* L.)

**DOI:** 10.1101/2024.10.30.621147

**Authors:** Saad Farid Usmani, Muhammad Abu Bakar Saddique, Muhammad Hammad Nadeem Tahir, Hafiz Nazar Faried

## Abstract

Chili (Capsicum annuum L.) also known as hot pepper and red pepper is a valuable condiment all around the world. It holds significant importance in the global economy and is used in a variety of products such as sauces, pickles, medicines, and insect-repellent sprays. Pakistan is 4^th^ largest chili producer worldwide. Global warming a is an important issue around the globe that causes losses to global agricultural productivity. In this study, chili germplasm was screened to identify heat-tolerant genotypes. The experiment was carried out following factorial under RCBD (randomized complete block design) having 785 with two treatments and three repeats. The genotypes were sown in nursery trays in November 2023 and transplanted in the field in February 2024. The data recording was performed twice. The data collected in March-April was considered as data from controlled treatment while data recorded during June-July was considered as heat-stressed treatment. The data was recorded for Days to flowering, NDVI value, plant height (cm), number of primary branches per plant, fruit length (cm), fruit diameter (cm), individual fruit weight (g), number of fruits per plant, fruit yield per plant, and pollen viability. Heat susceptibility indices for all traits were calculated. The data was subjected to analysis of variance, path coefficient, and biplot analysis. Genotypes D1, D4, D7, D12, 12, 42, 75, 76, 118, 172, 217, 229, 424, 497, 514, 532, and 772 are selected as heat tolerant genotypes because the yield of these genotypes was unaffected or slightly affected due to heat stress during the experiment, while on the other hand, the genotypes 41, 44, 57, 63, 94, 123, 141, 154, 285, 347, 516, 540, 601 and 663 are selected as heat susceptible because these genotypes have high heat susceptibility index and there was a huge decrease in the yield in heat-stressed condition as compared to non-stressed condition.

## 1. Introduction

Chili (*Capsicum annum*), also known as red pepper or hot pepper, belongs to the Solanaceae family. Chili has been a part of the human diet since ancient times and is grown worldwide. Archeologist studies show that chili was domesticated 6000 years ago and was one of the earliest self-pollinating crops in Central and South America ^1^. Pakistan is the sixth largest producer of chili and the area of Pakistan under the chili is 157.9 thousand acres with a production of 142.9 thousand tons ^2,3^. Globally, the average output is 1857 thousand hectares. Sindh is the largest producer of chili with more than 80% of total national chili production ^4^. Fresh and dried chili is essential to the daily diet in Pakistan as well as all around the world ^5^. Fresh chili acts as a healing agent and strengthens the immune system, especially in the event of cellular damage ^6^. Moreover, chili is used in medicine as an antibiotic.

The chili plant is susceptible to heat stress; elevated temperatures impact almost every stage of development. Floral aberration, Pollen viability, fruit set, and the number of seeds per fruit are all negatively impacted (Rosmaina *et al*., 2021). Additionally, it alters the fruit’s color and chemical makeup, lessening its flavor, and ultimately leading to low-yield and low-quality chilies ^9^. The optimum temperature for higher crop productivity of chili is between 22°C and 30 °C ^5^. High temperatures harm pepper plant health and yield although temperatures exceed 40 degrees Celsius during May and June.

Because of the adverse and unfavorable climatic conditions, climate change has become an important concern for agricultural productivity. Increasing crop productivity is challenging due to rising temperatures and decreased crop water availability brought on by climate change and global warming ^10^. The temperature is rising, which is extremely dangerous for crops and, in turn for food security. To minimize the negative impacts of climate change and global warming and to maintain chili productivity, it is necessary to select and produce heat-tolerant cultivars and to adopt sustainable farming practices ^11^.

The heat susceptibility index (HSI) shows us how much a plant is susceptible to heat stress and is commonly used for indirect selection to identify heat-responsive genotypes in plants ^12^. Plants with a low heat susceptibility index during high temperatures, tend to have better yield potential. The normalized difference vegetative index (NDVI) effectively indicates the stay-green trait, reflecting overall plant health. Selecting these traits in breeding can enhance reproductive success, photosynthesis, and other yield-related traits under heat stress. Pollen viability is also a key trait for selecting heat-tolerant genotypes. Pollen is the most heat susceptible tissue in many crops, and without viable pollen, the fruit set is reduced or completely impeded ^13^. As a plant breeder, it is crucial to choose those physiological traits of plants that help plants to survive in stressed conditions and also provide optimum yield in the harsh environment.

Conserving genetic resources in biodiversity and understanding the role of genes is crucial for both the species that possess this biodiversity and for humanity as a whole ^14^. This conservation is essential for future food security and maintaining biodiversity. In agriculture, a major concern about biodiversity loss is the reduction of genetic diversity, also known as “genetic erosion.” The genetic basis of many commercially cultivated species has narrowed genetic base due to intense yield improvement processes, limiting the number of genotypes available for breeding programs (Begna, 2021).

Most genetic variability in plants is found in their wild relatives, which still exist in centers of diversity ^16^. Wild relatives have always been valuable to breeders, who use hybridization to transfer genes from native and exotic species to cultivated ones, enhancing traits like stress tolerance or pest resistance. Genetic improvement in crops is a cost-effective, long-term, and sustainable way to address the harmful effects of abiotic stresses. Developing climate-resilient cultivars is crucial for improving chili and other crops. Utilizing the genetic variation in chili germplasm for heat tolerance is a promising approach.

## 2 Materials and Methods

### 2.1 Plant Material

The chili germplasm was collected from the National Agricultural Research Center (NARC), Islamabad Pakistan, World Vegetable Center, Taiwan, and imported from the online European international market. A total of 785 genotypes including 12 demonstration genotypes (D1 to D12) were used in the experiment to identify heat-tolerant genotypes.

### 2.2 Experimental site

The field experiment was conducted over 10 months (Nov 2023 to Aug 2024) on the Experimental Farm of the MNS University of Agriculture (MNSUAM) Multan, Pakistan (30°8″ N, 71°26″ E and 215 m above sea level). The climate in this region is subtropical with an average annual rainfall of 175 mm, 90% of which falls from July to September. The monthly average maximum temperature ranges from 21 °C in January to 38 °C in April, while the monthly average maximum temperature ranges from 32 °C to 46 °C in June. During the trial period (November 2023 to Aug 2024), monthly mean maximum and minimum temperatures were recorded and represented.

### 2.3 Layout and experimental conditions

The experiment followed factorial under a randomized complete block design (RCBD) with two treatments and three repeats. The seedlings were grown in seedling trays filled with peat moss in the greenhouse, under controlled conditions to get optimum germination percentage and a healthy plant nursery. 45-day-old seedlings were transplanted in the research area of MNS University of Agriculture, Multan (MNSUAM). Transplanting was done by hand after proper land preparation at a distance of 45 cm between plants and 75 cm between rows. Data recording was performed twice, i.e. data recorded during April-May 2024 was considered controlled data, and data recorded during June-July 2024 was considered heat-stressed data ^7^. Three plants per replication were used to record data in both treatments.

### 2.3 Data collection for morpho-physiological traits

#### 2.3.1 Germination Percentage (%)

The germination percentage was calculated by the following formula.

Germination percentage (%) = (No of seeds germinated / Total number of seeds) *100

#### 2.3.2 Days to 50% flowering (DAT)

The number of days after transplant (DAT) was calculated when 50% of the total plants, initiated flowering.

#### 2.3.3 NDVI value

NDVI value was estimated with the help of NDVI-CM1000 chlorophyll meter during sunshine between 7 am-10 am. This meter estimates the NDVI value by the given formula NDVI = (NIR - Red) / (NIR + Red)

(NIR is near-infrared light and Red is visible red light).

#### 2.3.4 Plant Height (cm)

Data for plant height of three plants from each genotype in each replication were collected from the soil surface to shoot tip by measuring tape.

#### 2.3.5 Number of primary branches

The number of branches that originate from the main stem were counted as primary branches.

#### 2.3.6 Pollen viability (%)

Newly opened flower buds were collected and fixed in Carnoy’s fixative. Later, Alexander staining was used to check pollen viability (%). Stained pollens were examined under the microscope and viable pollens were observed. Pollen viability percentage was estimated by the given formulae

Pollen viability % = (Viable pollens / total pollen) *100

#### 2.3.7 Fruit length (cm)

The length of fruits was measured by using a digital vernier caliper.

#### 2.3.8 Fruit diameter (cm)

A digital vernier caliper was used to measure the fruit diameter of each genotype from each replication.

#### 2.3.9 Individual Fruit Weight (g)

Digital weight balance was used to measure the individual fruit weight. Data of fruits from each replication were collected and their average was computed.

#### 2.3.10 No of fruits per plant

The ripe fruits of tagged three plants were harvested, counted, and summed on each picking, and their average data were collected.

#### 2.3.11 Fruit yield per plant (g)

The ripe fruits of the three best-performing plants selected and tagged plants were harvested and weighed with digital weight balance in grams.

#### 2.3.12 Heat Susceptibility Index (HSI)

It was calculated by using formulae

HSI = (1-YD /YP)/D

Where, YD = Mean data of a genotype at high temperature

YP = Mean data of genotype at optimum temperature

D (stress intensity) = 1-(mean YD of all genotypes/mean YP of all genotypes)

#### 2.2.4 Statistical analysis

The collected data were analyzed by variance analysis (ANOVA) to find the variation among genotypes (Steel *et al*., 1997). The correlation analysis was performed to figure out the effect of one trait over the other trait ^18^, while to determine the direct and indirect relationships between numerous yield-inducing and heat-tolerant traits, path coefficient analysis was performed ^19^. Biplot analysis was performed to test the response of genotypes for yield and heat-tolerance traits ^20^.

### 2.3 Results

#### 2.3.1 Analysis of variance (ANOVA)

The ANOVA (Analysis of Variance) was performed through the software “R Studio” to find significant differences between the genotypes and treatments for their morpho-physiological traits. ANOVA shows that there is a significant difference at p≤ 0.01, and P≤0.001between genotypes and their treatments morpho-physiological traits **DF**: Days to Flowering, **NDVI**: Normalized difference vegetation index, **PH**: Plant Height, **NOB**: Number of primary branches, **FL**: Fruit length, **FD**: Fruit Diameter, **SFW**: Single fruit weight, **PV**: Pollen Viability, **YPP**: fruit yield per plant and **NOF**: No. of fruits per plant

#### 2.3.2. Clustered Heat-map Analysis

#### 2.3.3 Correlation and path co-efficient analysis

Assessing correlation coefficients helps plant breeders to develop breeding strategies for the evaluation of specific genotypes. The plant character association shows a relationship that might be different between the parents because of the different combinations of the source material and the environment in which the experiment has been performed.

Pearson correlation was performed to check degrees of associations between all morphological and physiological traits as given in Figure 2.6. Findings in this study suggest that the fruit diameter highly positively correlates with the single fruit weight. The fruit length also positively correlates with single fruit weight and fruit diameter. The yield of plants positively correlates with all other traits such as NDVI value, plant height, number of branches, fruit length, fruit width, single fruit weight, and pollen viability. Plant height has a strong positive correlation with the number of primary branches. Pollen viability (PV) shows a positive correlation with yield per plant and number of fruits per plant but it has a non-significant relation with plant height and number of branches. Day to flowering shows a negative correlation with the number of fruits, yield per plant, single fruit weight, fruit length, and fruit diameter, however, shows a non-significant correlation with plant height, number of branches, and NDVI. This means yield-related traits increase with the decrease in number of days to flowering.

**Figure 2.1:**
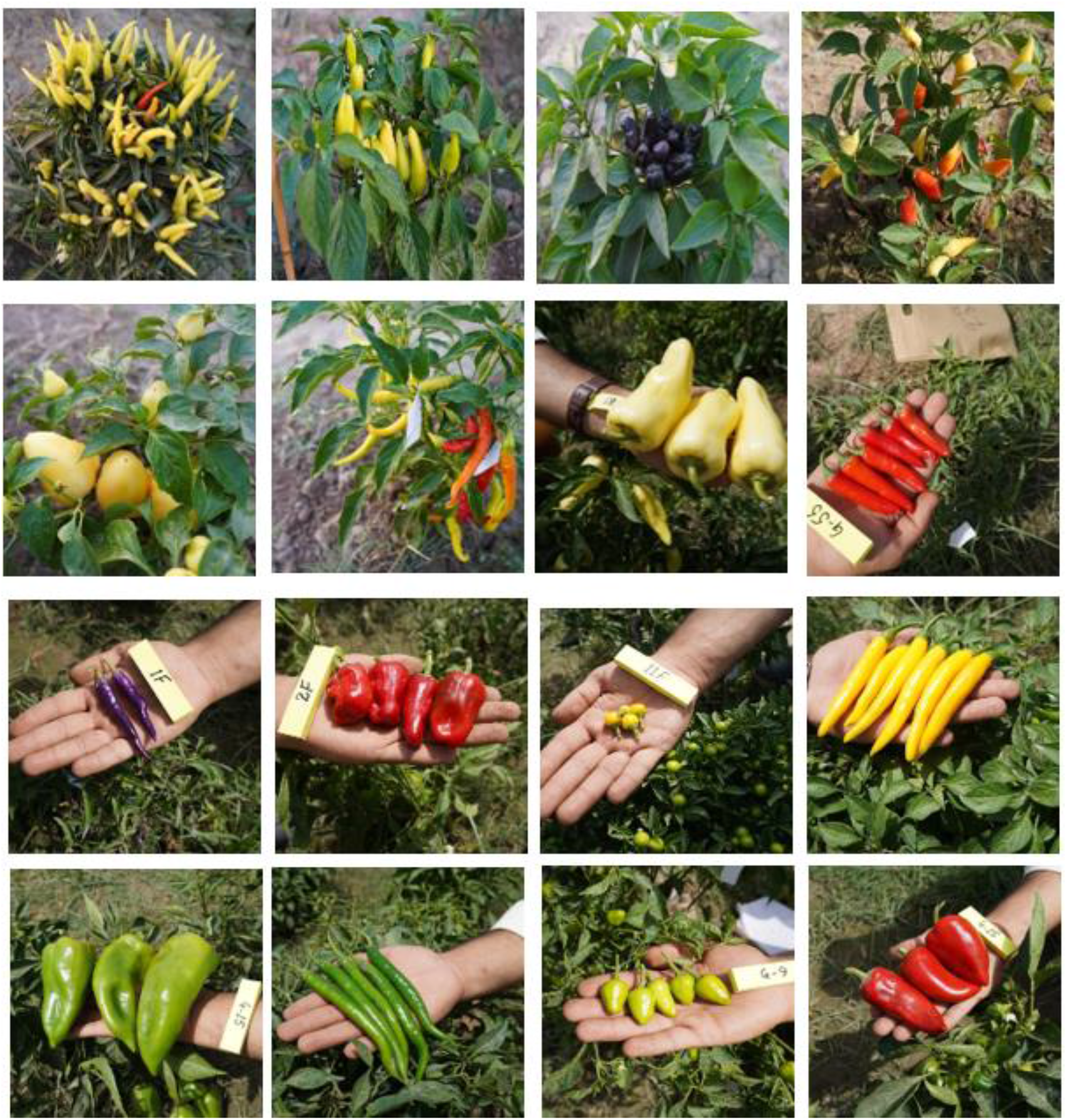
Different genotypes of chili heaving different fruit shapes and colors.

**Figure 2.2.**
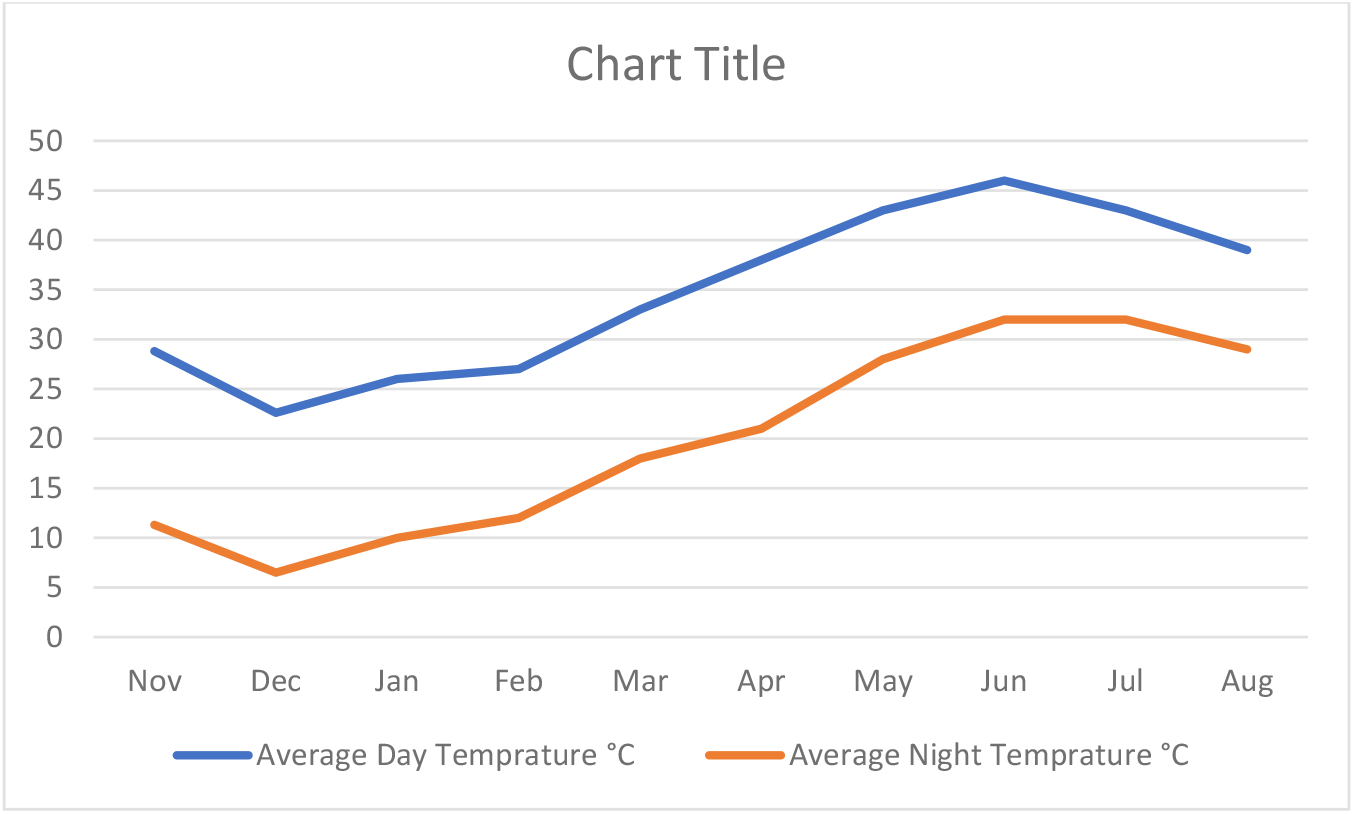
Average Day and Night Temperature (°C) from November 2023 to August 2024.

**Figure 2.3:**
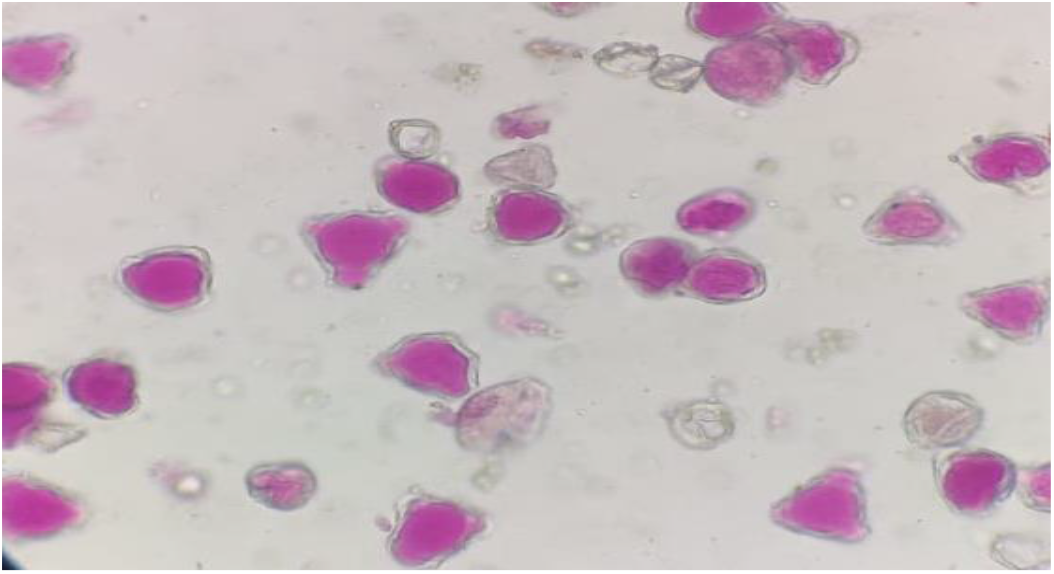
Microscopic view of chili pollens, stained for estimation of pollen viability

Path-coefficient analysis was performed to check direct and indirect relationships among morpho-physiological and yield traits of chili genotypes ^19^. Path analysis was performed using “R Studio” as shown in Figure 2.6. The arrows in the path analysis show the direct relationship between those particular traits.

#### 2.3.4 Biplot analysis for selection of chili heat tolerant and heat susceptible genotypes

This experiment evaluated best-performing genotypes for morpho-physiological and yield-related traits like NDVI, FL, FD, SFW, PV, YPP, and NOF. Highly significant variations were identified for all the traits (p<0.01, table 2.3). The biplot analysis indicates the performance of genotypes for yield and quality-related parameters. Using Biplot analysis, genotypes were selected based on heat response and morphological parameters. This approach involves distributing all genotypes in the proper environment based on the attributes under study. Figure 2.4 illustrates a combined biplot comparing the effectiveness of each genotype under non-stress and heat-stress conditions. Based on the biplot selection of genotypes, heat tolerance and average best performance on all mentioned morphological traits were performed by keeping in view the origin and line (OP) distance and the region in which genotypes were present.

**Table 2.2:**
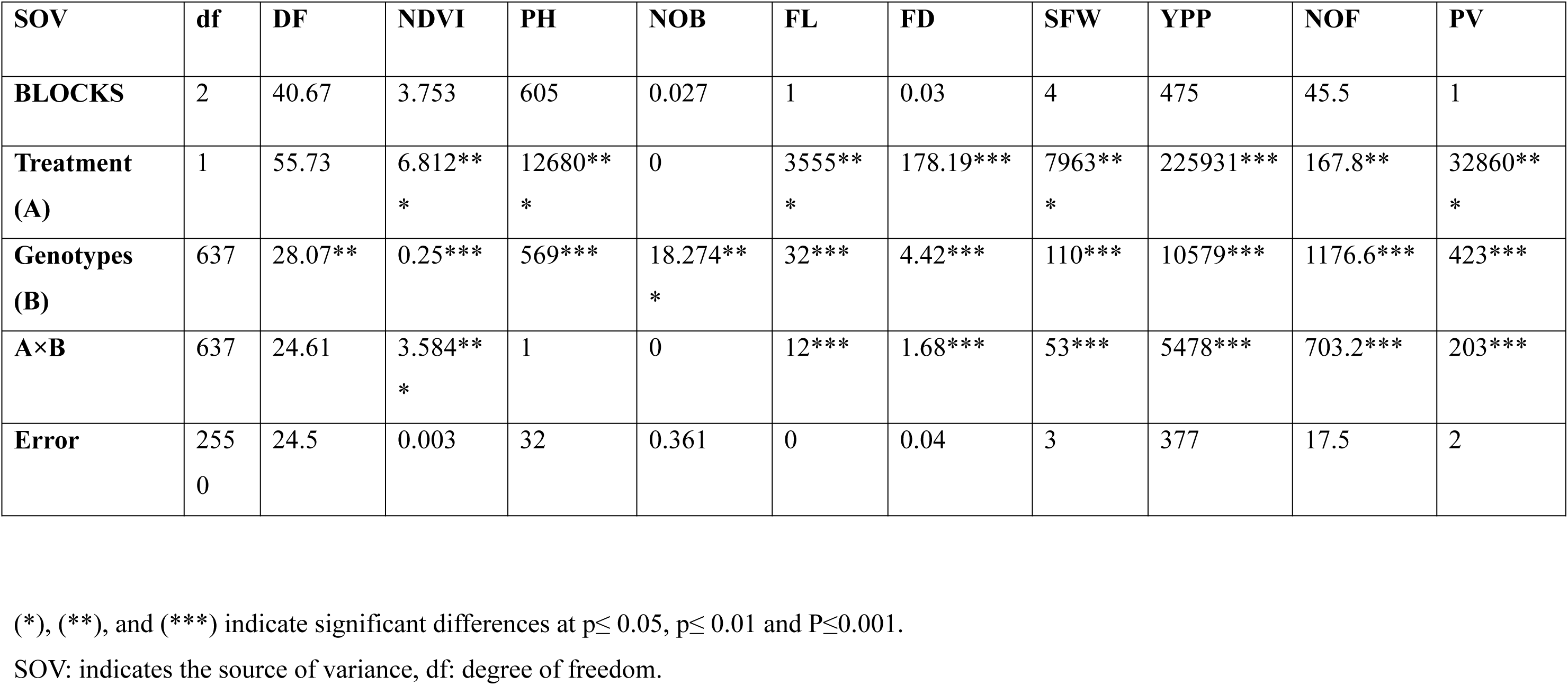
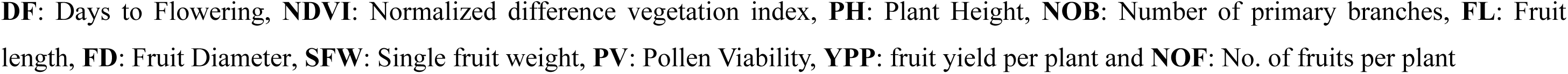
Mean square values of morpho-physiological traits of chili.

**Table 2.4:** Heat Susceptibility Indices of Yield-related Traits for heat tolerant chili genotypes

| **Genotype** | **FL** | **FD** | **SFW** | **YPP** | **NOF** | **PV** |
| --- | --- | --- | --- | --- | --- | --- |
| D1 | 0.17 | 0.00 | 0.38 | -2.87 | -13.40 | 0.94 |
| D4 | 0.42 | -0.98 | 0.12 | 0.12 | -1.78 | 0.94 |
| D7 | 0.38 | -0.91 | 0.22 | -0.64 | -3.08 | 1.06 |
| D12 | 0.58 | 0.54 | 0.38 | -1.41 | -14.78 | 0.82 |
| Big Red Thick | 0.70 | -0.46 | 0.31 | -4.23 | -42.44 | 0.45 |
| Red Pointed | 0.64 | -0.06 | 0.57 | -1.77 | -14.86 | 0.72 |
| AVPP0303 | -0.05 | -1.10 | 0.92 | -7.89 | -143.89 | 0.82 |
| AVPP0206 | 0.32 | 0.52 | 0.00 | -2.14 | -29.52 | 0.90 |
| VI012902 | 0.19 | -0.29 | 0.47 | -5.66 | -69.17 | 0.89 |
| VI012005 | 0.32 | 1.00 | 0.61 | -1.14 | -31.03 | 1.04 |
| VI011992 | 0.92 | 0.77 | 1.15 | -1.00 | -90.24 | 0.94 |
| 39553 | 0.30 | 0.06 | 0.72 | -7.66 | -121.30 | 0.58 |
| VI037513 | 0.19 | -0.55 | 0.72 | -9.44 | -113.92 | 0.73 |
| 32319 | 0.74 | 0.45 | 0.82 | 0.24 | -15.31 | 1.06 |
| Aji champion | 0.05 | 0.04 | 0.65 | 0.44 | -1.23 | 0.91 |
| Golden cayenne | 0.47 | 1.37 | 0.55 | -2.79 | -28.00 | 1.04 |
| Sibirischer<br>Hauspaprika | 0.12 | 0.39 | 0.00 | -3.38 | -33.68 | 0.93 |
| Jalaro | 0.13 | 0.88 | 0.76 | -0.57 | -5.00 | 0.80 |
| Botinecka Zuta | 0.43 | -0.02 | 1.05 | -1.98 | -39.11 | 0.91 |
| Nocturne | 0.95 | 0.57 | 1.01 | 0.29 | -11.36 | 1.34 |
| AVPP0310 | 0.65 | -0.25 | 0.85 | 0.64 | -15.16 | 1.03 |
| AVPP0516 | -0.05 | 0.17 | 0.47 | -8.50 | -123.94 | 0.91 |
| **D-Values** | **0.50** | **0.36** | **0.65** | **0.41** | **0.05** | **0.09** |

**Table 2.5:** Heat Susceptibility Indices of Yield-related Traits for heat Susceptible chili genotypes

| Genotype | FL | FD | SFW | YPP | NOF | PV |
| --- | --- | --- | --- | --- | --- | --- |
| Red Shine | 2.02 | 2.74 | 1.53 | 2.43 | 20.00 | 1.17 |
| yellow sunshine | 2.02 | 2.74 | 1.53 | 2.43 | 20.00 | 1.17 |
| yellow long | 2.02 | 2.74 | 1.53 | 2.43 | 20.00 | 1.06 |
| round red pointed | 2.02 | 2.74 | 1.53 | 2.43 | 20.00 | 1.17 |
| VI012893 | 2.02 | 2.74 | 1.53 | 2.43 | 20.00 | 0.82 |
| VI011989 | 2.02 | 2.74 | 1.53 | 2.43 | 20.00 | 0.94 |
| VI056423 | 2.02 | 2.74 | 1.53 | 2.43 | 20.00 | 0.82 |
| VI037629 | 2.02 | 2.74 | 1.53 | 2.43 | 20.00 | 0.94 |
| VI012520 | 0.97 | 0.54 | 0.71 | 2.23 | 17.27 | 1.17 |
| VI012121 | 0.00 | 0.53 | 0.87 | 2.05 | 14.00 | 0.82 |
| Hungarian-hot | 1.17 | 1.14 | 1.18 | 2.10 | 11.79 | 1.06 |
| Thai Orange Hot | 0.58 | 1.16 | 1.05 | 2.03 | 12.92 | 0.82 |
| Satan's Kiss | 0.62 | 0.56 | 1.11 | 1.99 | 10.69 | 0.94 |
| Marconi Red | 1.06 | 0.82 | 1.07 | 1.95 | 11.67 | 1.06 |
| **D-Values** | **0.50** | **0.36** | **0.65** | **0.41** | **0.05** | **0.09** |

**Figure 2.4:**
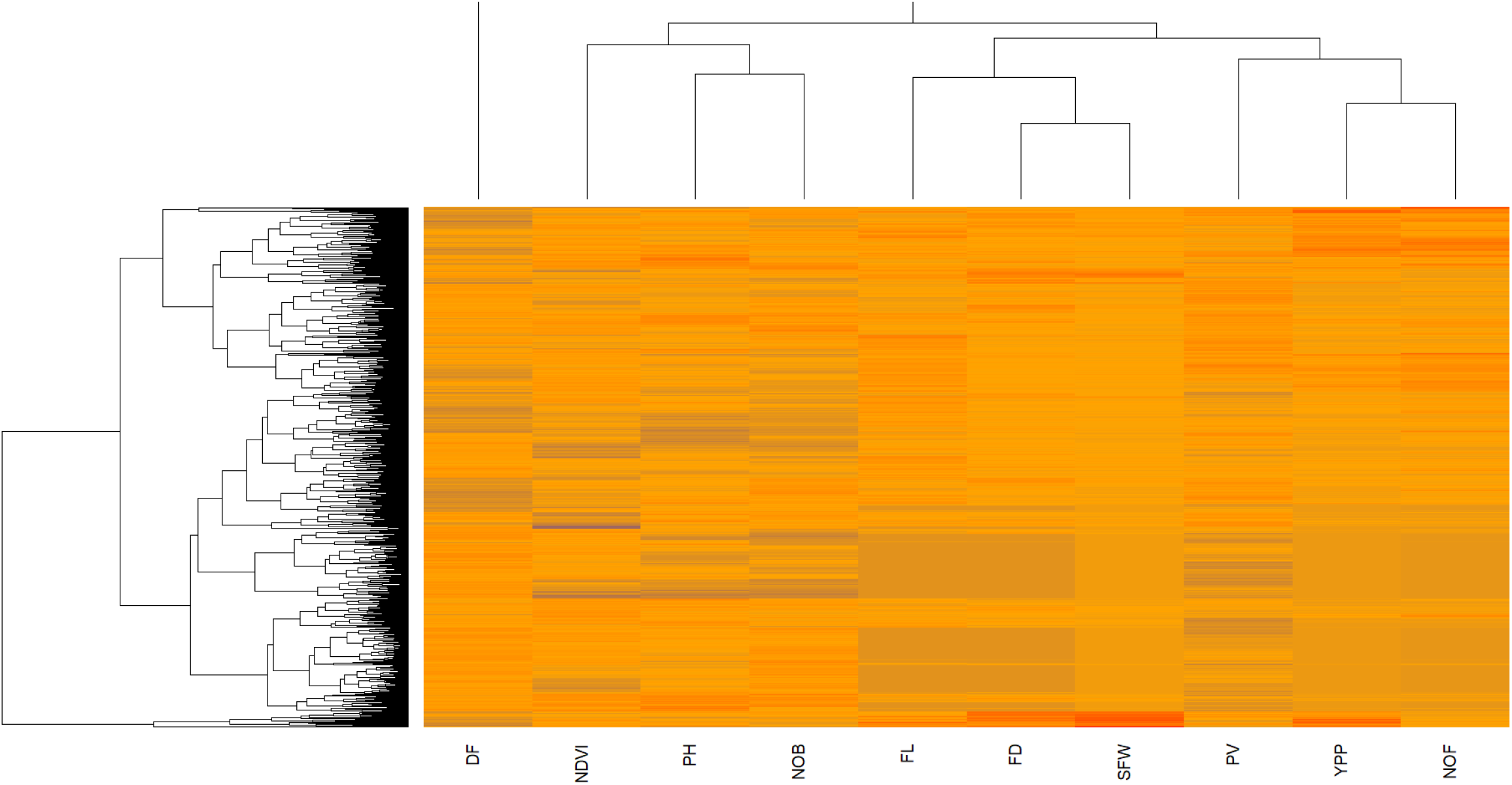
Heatmap and Dendrogram analysis of morpho-physiological and yield-related traits of Chili under non-stressed condition. The red color indicates the high magnitude the blue color indicates the low magnitude of the specific traits, and the dendrogram shows linkage.

**Figure 2.5:**
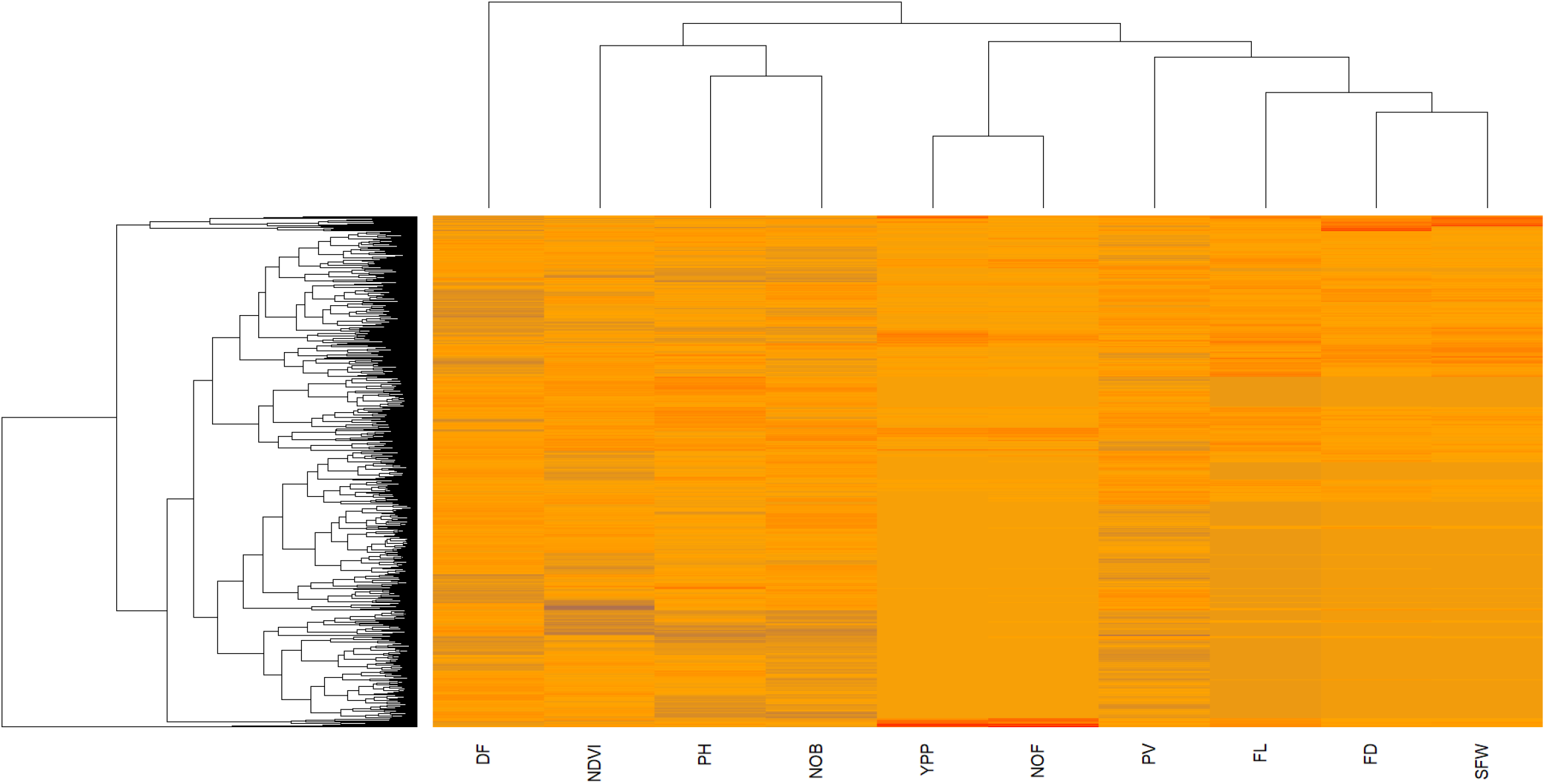
Clustered heat-map analysis of morpho-physiological and yield-related traits of Chili under Heat-stressed condition. Colors in the heatmap show the magnitude of values of different traits and the dendrogram shows the association within traits.

**Figure 2.6:**
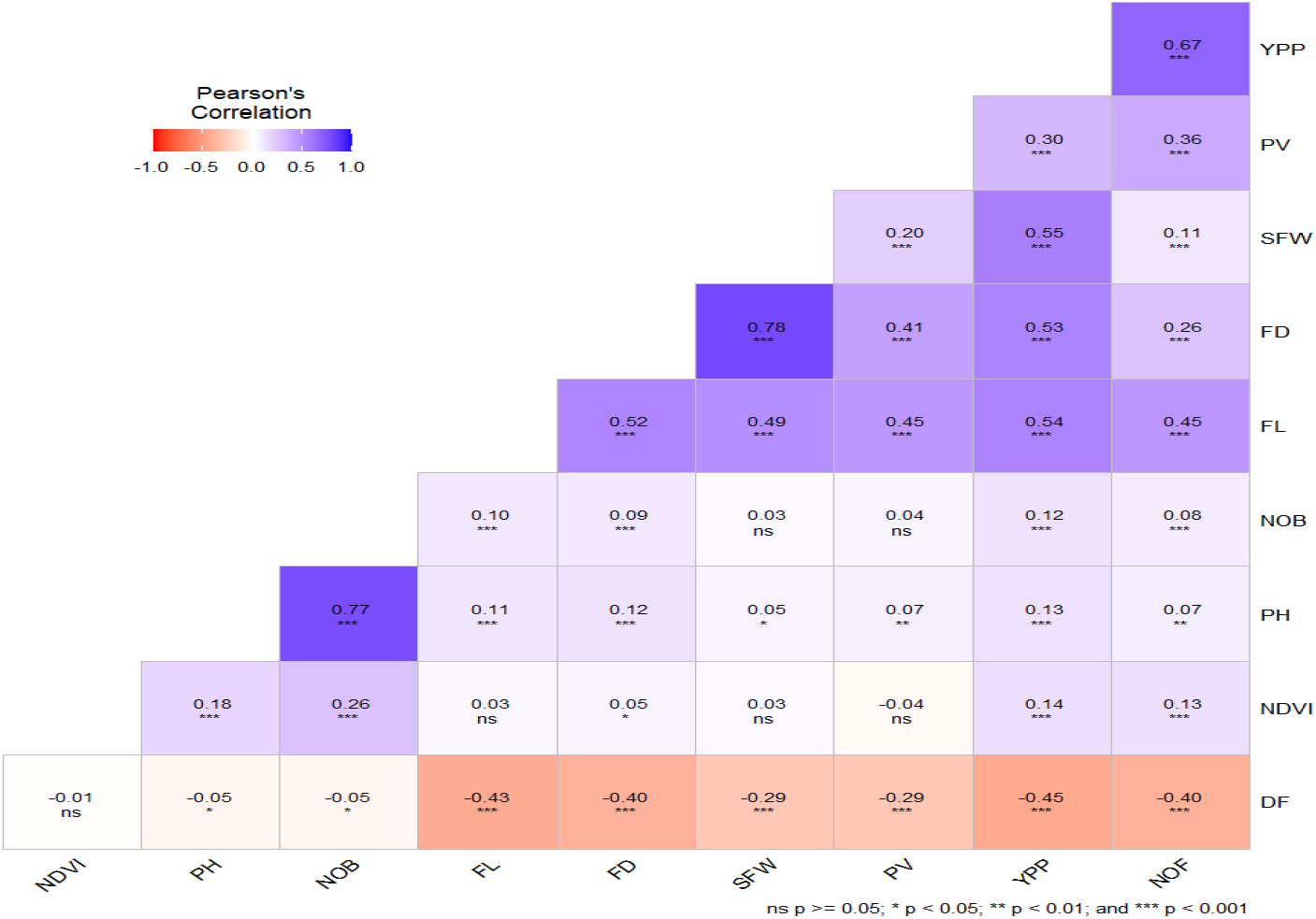
Correlation analysis under non-stressed conditions

**Figure 2.7:**
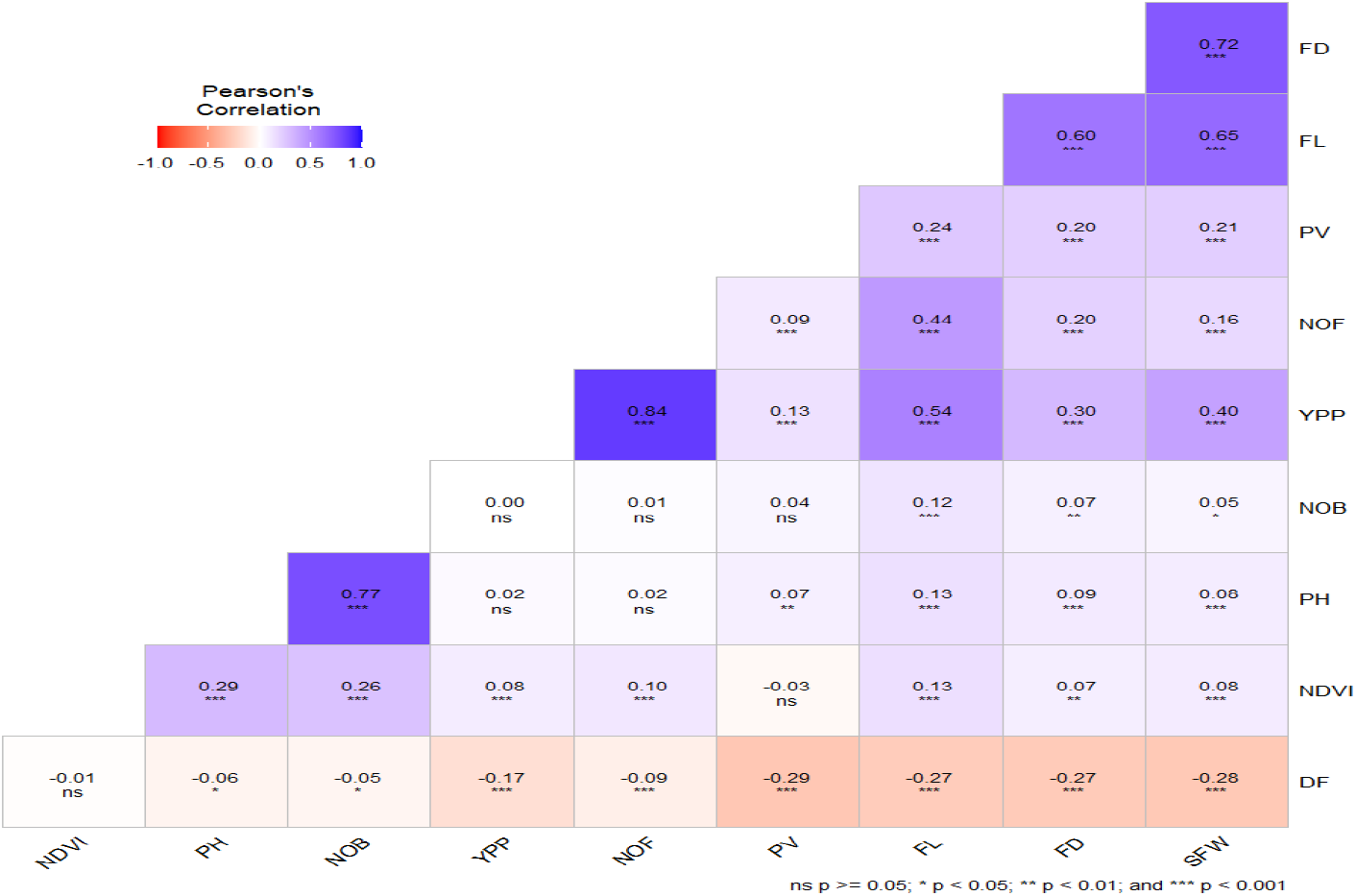
Correlation analysis under heat-stressed conditions. The blue color indicates the positive correlation, red color indicates the negative correlation between traits, however, (*), (**), and (***) indicate the significance level of correlation <0.05, <0.01, and <0.001

**Figure 2.8.**
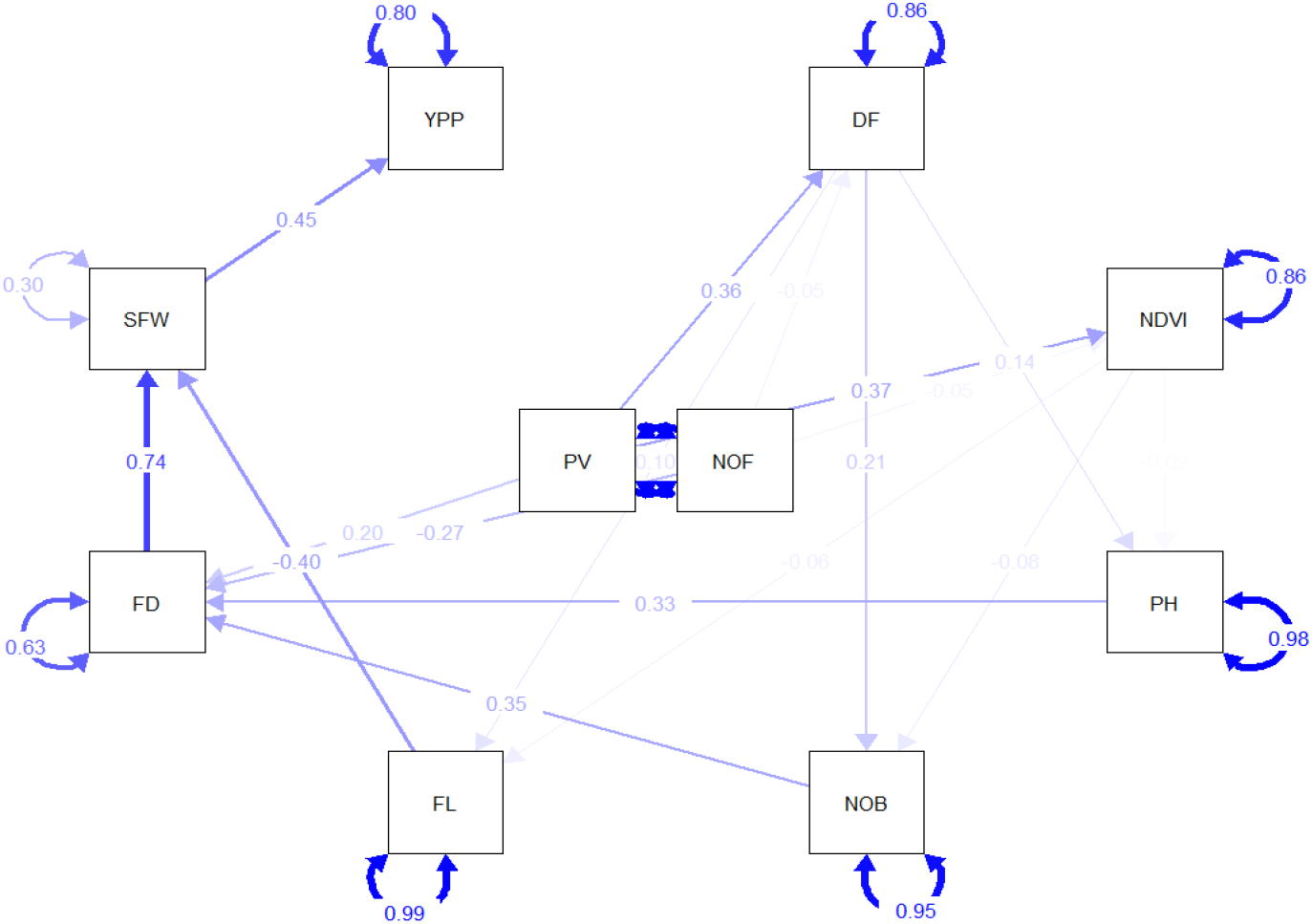
Path coefficient analysis under non-stressed conditions.

**Figure 2.9.**
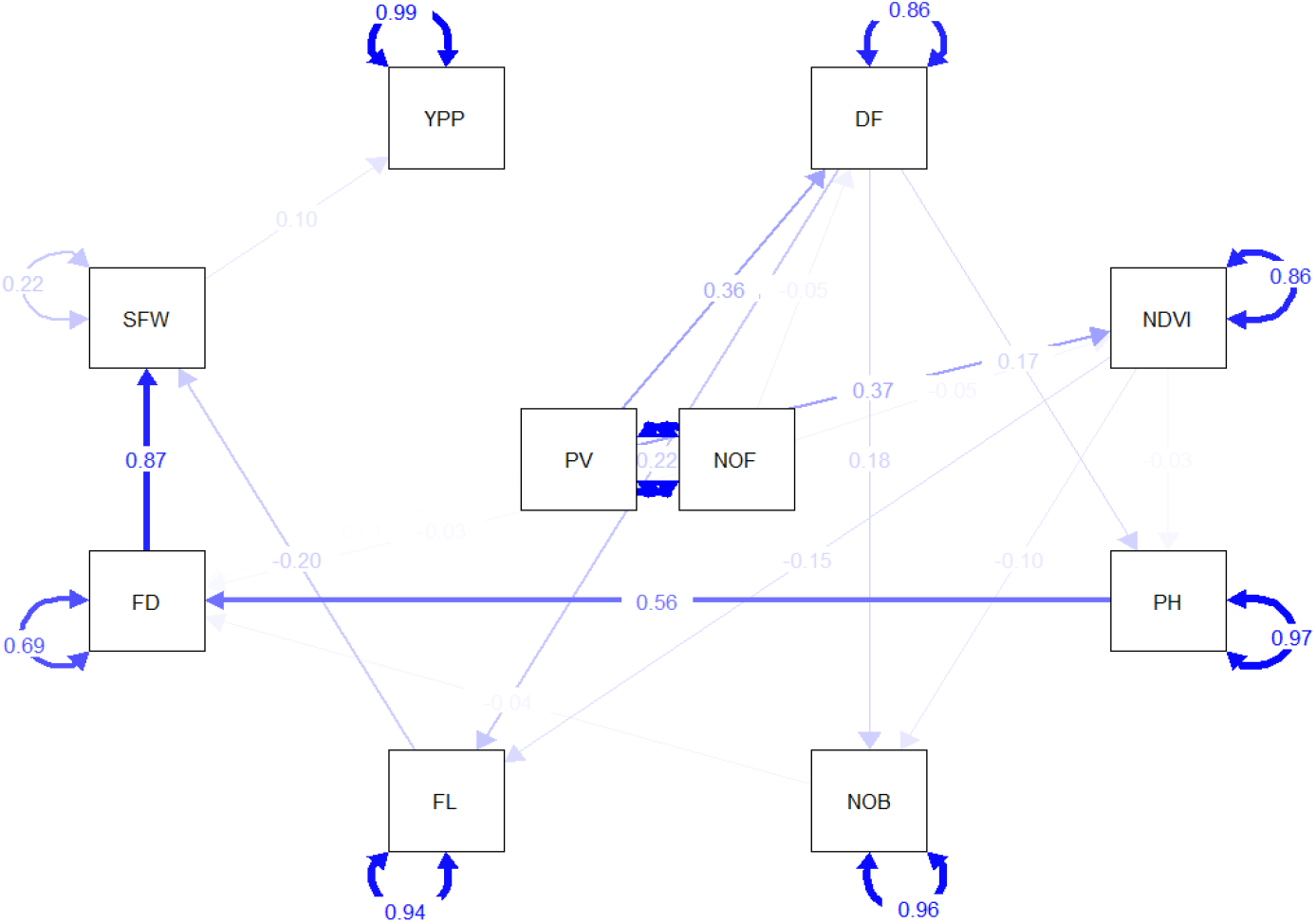
Path coefficient analysis under heat-stressed conditions.

**Figure 2.10:**
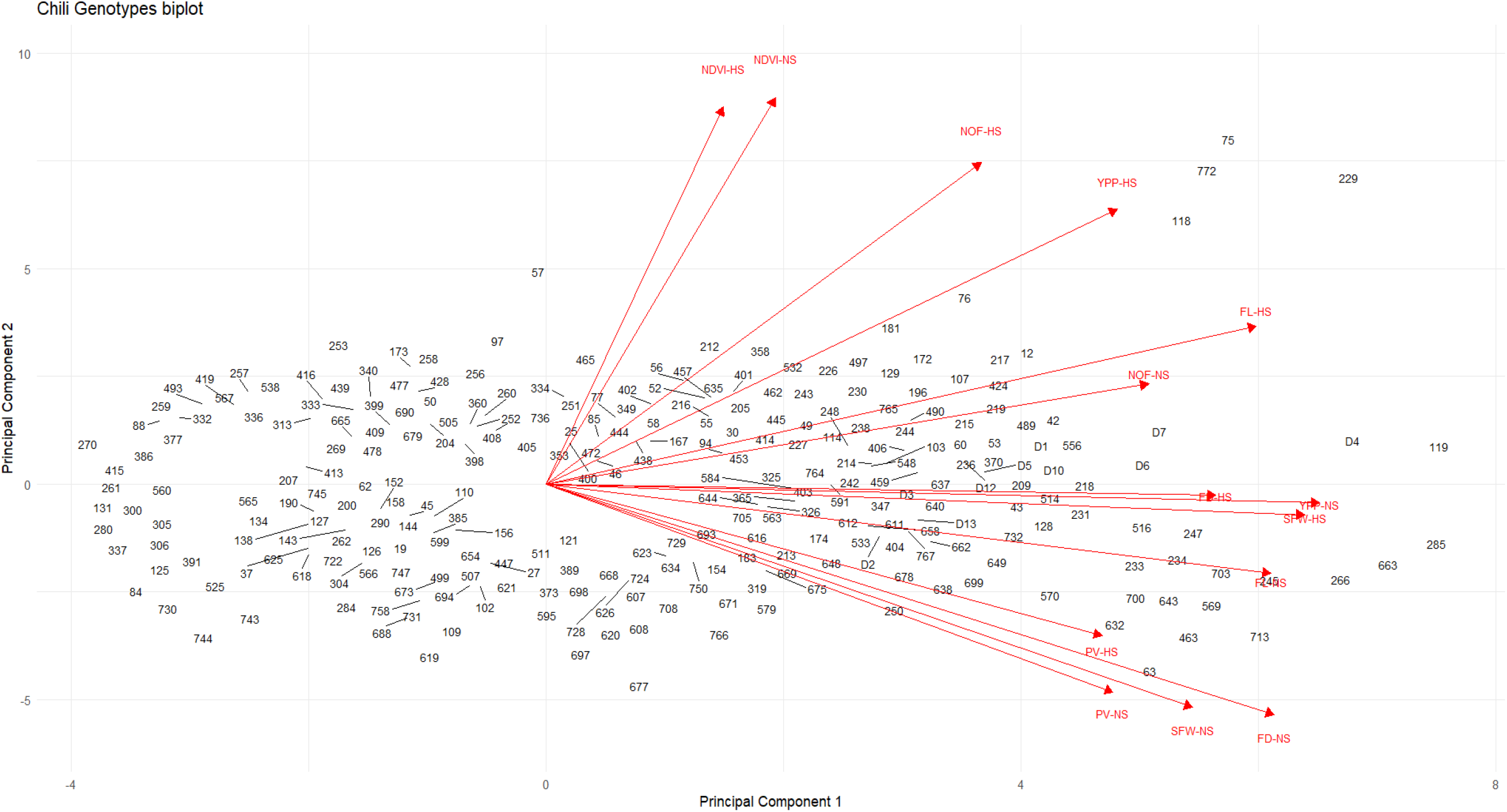
Biplot analysis of yield-related traits under non-stressed (NS) and heat-stressed (HS) conditions in chili

The genotypes D1, D4, D7, D12, 12 (Big Red Thick), 42 (Red Pointed), 75 (AVPP0303), 76 (AVPP0206), 118 (VI012902), 172 (VI012005), 217 (VI011992), 229 (39553), 424 (32319), 490 (Aji champion), 497 (Golden cayenne), 577 (Chinesischer Runzelhaut Chili), 635 (Sibirischer Hauspaprika), 640 (Jalaro), 703 (Nocturne) and 772 (AVPP0516) are selected as heat tolerant genotypes while on the other hand, the genotypes 41 (Red Shine), 44 (yellow sunshine), 58 (yellow long), 63 (round red pointed), 94 (VI012893), 123 (VI011989), 141 (VI056423), 154 (VI037629), 285 (VI012520), 347 (VI012121), 516 (Hungarian hot), 540 (Thai Orange Hot), 601 (Satan’s Kiss) and 663 (Marconi Red) are selected as heat susceptible from the Figure 2.4.

The biplot shows the performance of genotypes concerning the traits. All the genotypes are scattered across their respective trait vectors. In the biplot graph, NDVI stands for Normalized Difference vegetative index, PV is Pollen viability, FL is for fruit length, FD is for fruit diameter, NOF is for the number of fruits per plant and YPP stands for overall yield per plant.

#### 2.3.5 Heat Susceptibility Indices for Yield Contributing Traits

The Heat Susceptibility Index (HSI) is a measurement that determines how susceptible a crop or plant is to heat stress. It helps in determining which crop varieties are more resistant to high temperatures. The heat stress index (HSI) is calculated by comparing a plant’s yield or other performance measures to those under optimal growing conditions. It was calculated by using formulae (Fischer and Maurer, 1978).

HSI = (1-YD /YP)/D

### 2.4 Conclusion

Yield and stress-responsive traits are important traits for plant vegetative growth and development. Yield is a complex trait that is the function of many yield-contributing parameters like the height of plants, number of branches, fruit length, fruit diameter, fruit weight, and no. of fruits per plant. The performance of 785 chili genotypes was evaluated to identify the variation for important agronomic traits in field conditions.

Highly significant variation was identified in the genotypes and their treatments for PH, FS, FC, TSS, FL, FD, NOF, IFW, FY, and PV (Table 2.3). The knowledge of the correlation between plant traits and yield is important while implementing a selection pattern with the target for enhanced yield. Path coefficient analysis showed the direct and indirect association between morpho-physiological and yield-related traits of chili. The correlation coefficient helps the plant breeder select an efficient plant trait in a crop breeding program (Maga *et al*., 2012). Correlation analysis provides us with information about the selection of traits, which leads to enhancement for all positively correlated traits. Several plant traits are correlated due to association, this association can be positive or negative, with other traits (Wali and Kabura, 2014).

The biplot analysis indicates the performance of genotypes for yield and quality-related parameters. The distance of genotypes from their origin, high-yielding, and better-quality genotypes were selected based on their trait responses. Genotypes D1, D4, D7, D12, 12, 29, 42, 75, 76, 118, 217, 229, 424, 497, 514, 532, and 772 are selected as heat tolerant genotypes because the yield of these genotypes was unaffected or slightly affected due to heat stress during the experiment, while on the other hand, the genotypes 41, 44, 57, 63, 94, 123, 141, 154, 285, 347, 516, 540, 601 and 663 are selected as heat susceptible because these genotypes have high heat susceptibility index and there was a huge decrease in the yield in heat-stressed condition as compared to non-stressed condition.

## REFERENCES

1. Brown CH, Clement CR, Epps P, Luedeling E, Wichmann S. The paleobiolinguistics of domesticated chili pepper (Capsicum spp.). Ethnobiology letters. 2013;4:1–11.

2. Zahra N, Naeem N, Butt AI, Saeed MK, Akram J, Gulzar E. Determination of Aflatoxins in Different Varieties of Chillies Collected from Lahore, Pakistan. International Journal of Food Science and Agriculture. 2022;6(3):349–354.

3. Rais MUN, Mangan T, Sahito JGM, Qureshi NA. A trend analysis: Forecasting growth performance of production and export of chilli in Pakistan. Sarhad Journal of Agriculture. 2021;37(1):220–225.

4. Akhund S, Akram A, Hanif NQ, Qureshi R, Naz F, Nayyar BG. Pre-harvest aflatoxins and Aspergillus flavus contamination in variable germplasms of red chillies from Kunri, Pakistan. Mycotoxin research. 2017;33:147–155.

5. Idrees S, Hanif MA, Ayub MA, Hanif A, Ansari TM. Chili pepper. In: Medicinal Plants of South Asia. Elsevier; 2020. p. 113–124.

6. Garnier A, Shahidi F. Spices and herbs as immune enhancers and anti-inflammatory agents: A review. Journal of Food Bioactives. 2021;14.

7. Kaur N, Dhaliwal MS, Jindal S, Singh P. Evaluation of hot pepper (Capsicum annuum L.) genotypes for heat tolerance during reproductive phase. International Journal of Bioresource and Stress Management. 2016;7(Feb, 1):126–129.

8. Rosmaina R, Utami D, Aryanti E, Zulfahmi Z. Impact of heat stress on germination and seedling growth of chili pepper (Capsicum annuum L.). IOP Conference Series: Earth and Environmental Science. 2021;637:12032. doi:10.1088/1755-1315/637/1/012032

9. Khaitov B, Umurzokov M, Cho K-M, Lee Y-J, Park KW, Sung J. Importance and production of chilli pepper; heat tolerance and efficient nutrient use under climate change conditions. Korean Journal of Agricultural Science. 2019;46(4):769–779.

10. Habib-ur-Rahman M, Ahmad A, Raza A, Hasnain MU, Alharby HF, Alzahrani YM, Bamagoos AA, Hakeem KR, Ahmad S, Nasim W. Impact of climate change on production; Issues, challenges, and opportunities in Asia. Frontiers in Plant Science. 2022;13:925548.

11. Uebersax MA, Cichy KA, Gomez FE, Porch TG, Heitholt J, Osorno JM, Kamfwa K, Snapp SS, Bales S. Dry beans (Phaseolus vulgaris L.) as a vital component of sustainable agriculture and food security—A review. Legume science. 2023;5(1):e155.

12. Fischer RA, Maurer R. Drought resistance in spring wheat cultivars. I. Grain yield responses. Australian Journal of Agricultural Research. 1978;29(5):897–912.

13. Giorno F, Wolters-Arts M, Mariani C, Rieu I. Ensuring reproduction at high temperatures: the heat stress response during anther and pollen development. Plants. 2013;2(3):489–506.

14. Salgotra RK, Chauhan BS. Genetic diversity, conservation, and utilization of plant genetic resources. Genes. 2023;14(1):174.

15. Begna T. Role and economic importance of crop genetic diversity in food security. International Journal of Agricultural Science and Food Technology. 2021 Apr 17:164–169. doi:10.17352/2455-815X.000104

16. Ebert AW, Engels JMM. Plant biodiversity and genetic resources matter! Plants. 2020;9(12):1706.

17. Steel RGD, Torrie JH. Principles and procedures of statistics. Principles and procedures of statistics. 1960.

18. Kwon SH, Torrie JH. Heritability of and interrelationships among traits of two soybean populations 1. Crop science. 1964;4(2):196–198.

19. Dewey DR, Lu K. A correlation and path-coefficient analysis of components of crested wheatgrass seed production 1. Agronomy journal. 1959;51(9):515–518.

20. Yan W, Cornelius PL, Crossa J, Hunt LA. Two types of GGE biplots for analyzing multi-environment trial data. Crop Science. 2001;41(3):656–663.

